## Supplementary Information for "Alternative architecture of the *E. coli* chemosensory array"

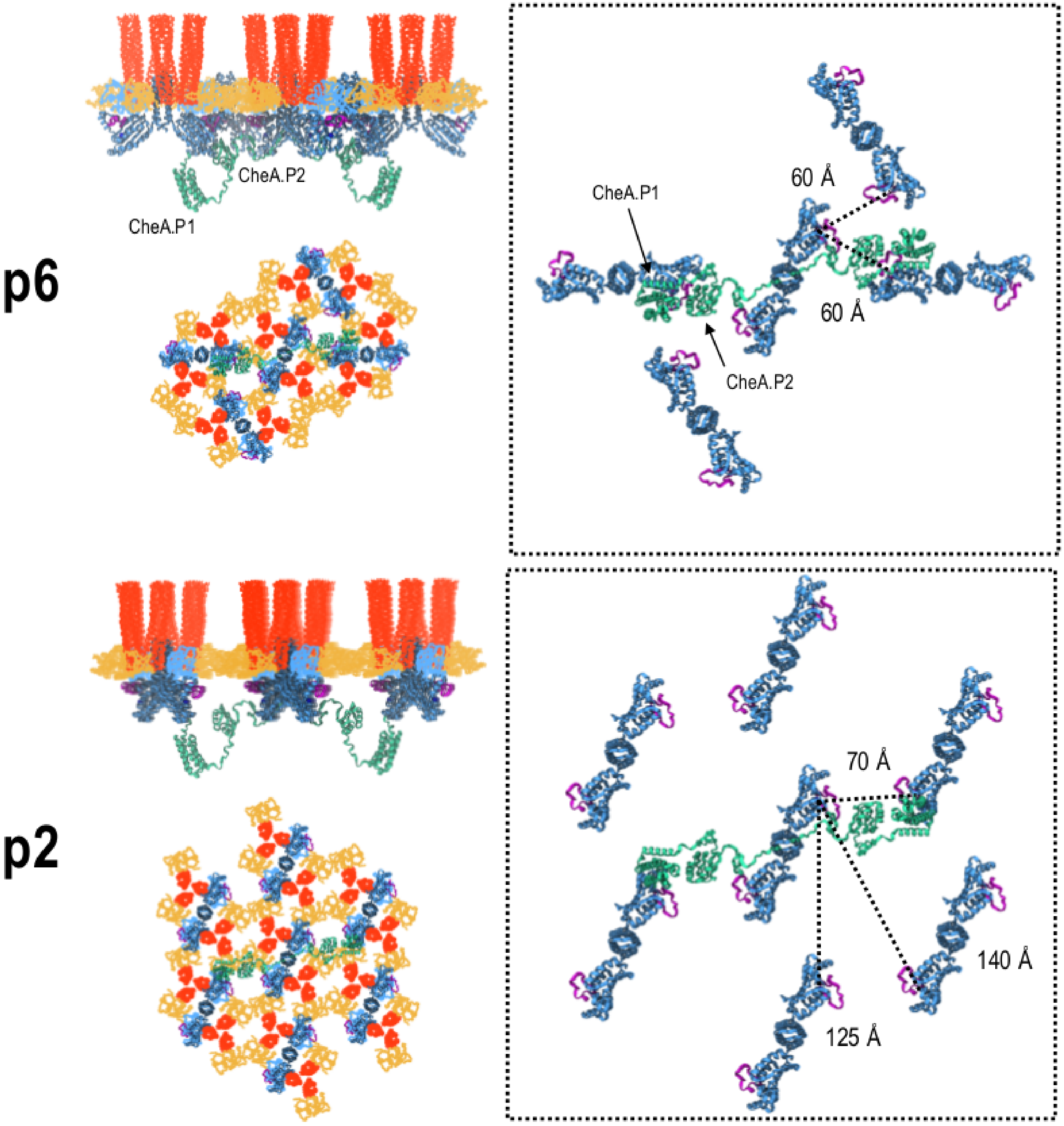


**Supplementary Figure 1. The intermolecular distance and the relative orientation of neighbouring CheA molecules are different between the p6 and p2 array architectures.** Side and bottom views of the p6- and p2-symmetric array models with the CheA.P1 and CheA.P2 domains (shown in green) modelled for a centralised CheA dimer. Receptors, CheA.P3.P4.P5, and CheW are coloured as in other figures. Boxed regions show the rough distances between an ATP-binding site from one of the central CheA monomers and ATP-binding sites on neighbouring CheAs that may be accessible in a trans-fashion by the CheA.P1 and CheA.P2 domains from the opposing central CheA monomer. The ATP lid (residues 455-475) of each CheA monomer is shown in purple. Receptors, CheW, and CheA.P5 are not shown for clarity.

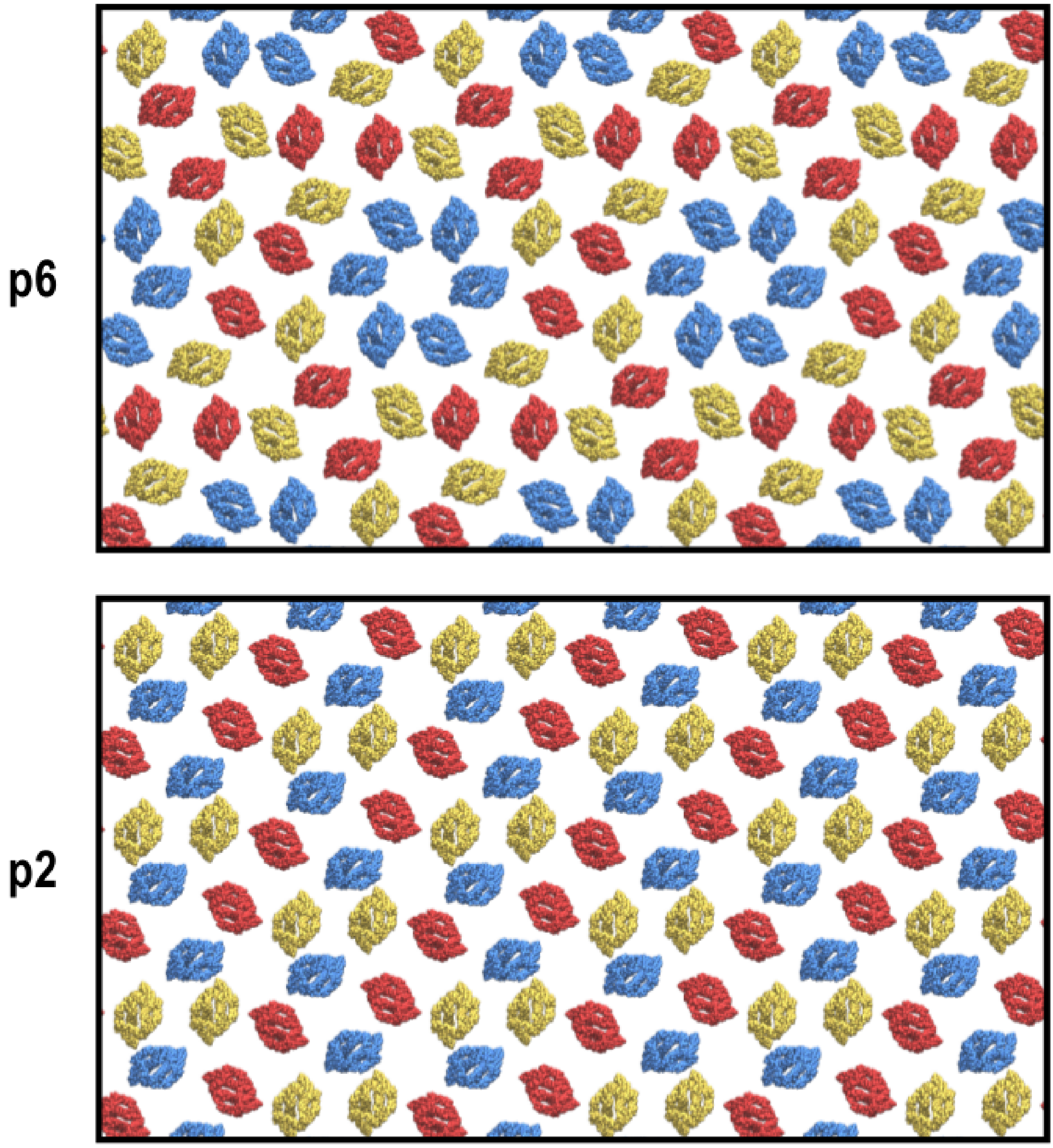


**Supplementary Figure 2. The hexagonal arrangement of receptors is conserved between the p6 and p2 array architectures.** Top view of the p6- and p2-symmetric array models with receptor periplasmic domains coloured according to baseplate partner. Receptors bound to CheA.P5 are shown in red, core CheW in yellow, and flanking CheW in blue.

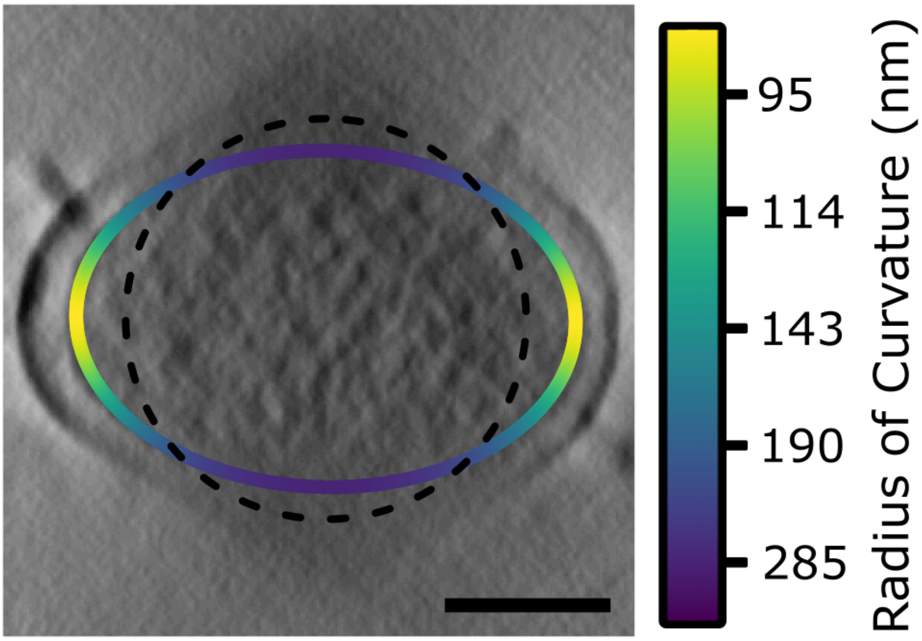


**Supplementary Figure 3. Estimation of the average radius of curvature of the inner membrane of the WM4196 minicells in cryo-ET.** An XZ projection through the region of a tomogram containing a p2-symmetric array architecture. Local estimates for radii of curvature are indicated on an overlaid ellipse. A circle with curvature equal to the mean curvature of the ellipse is plotted as a dashed line.

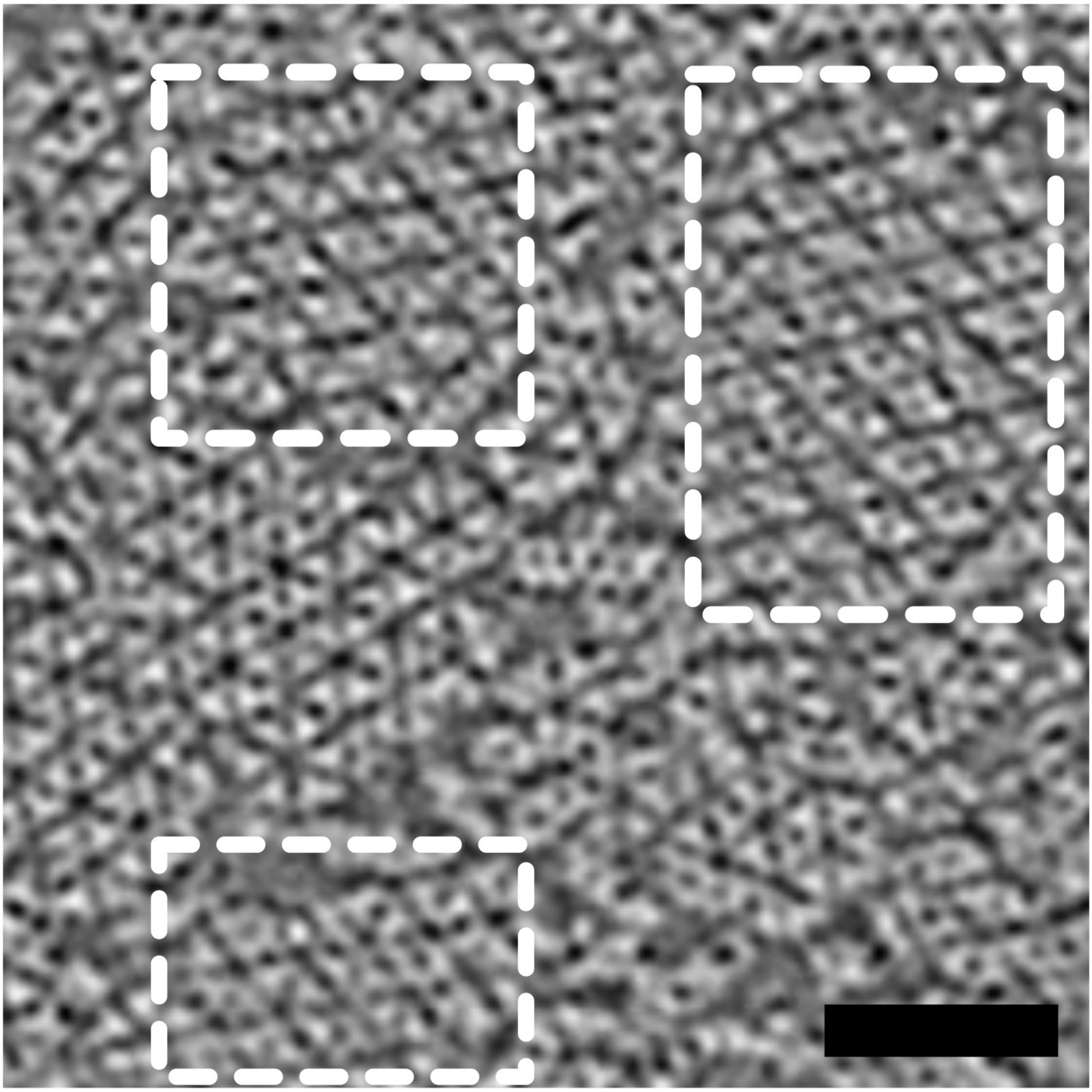

**Supplementary Figure 4. An image region of an *in vitro* reconstituted chemosensory array with very low curvature taken from Cassidy et al. 2020.** Parts on the image which appear to display a p2 array architecture are highlighted in dashed rectangles. Scale 50 nm.

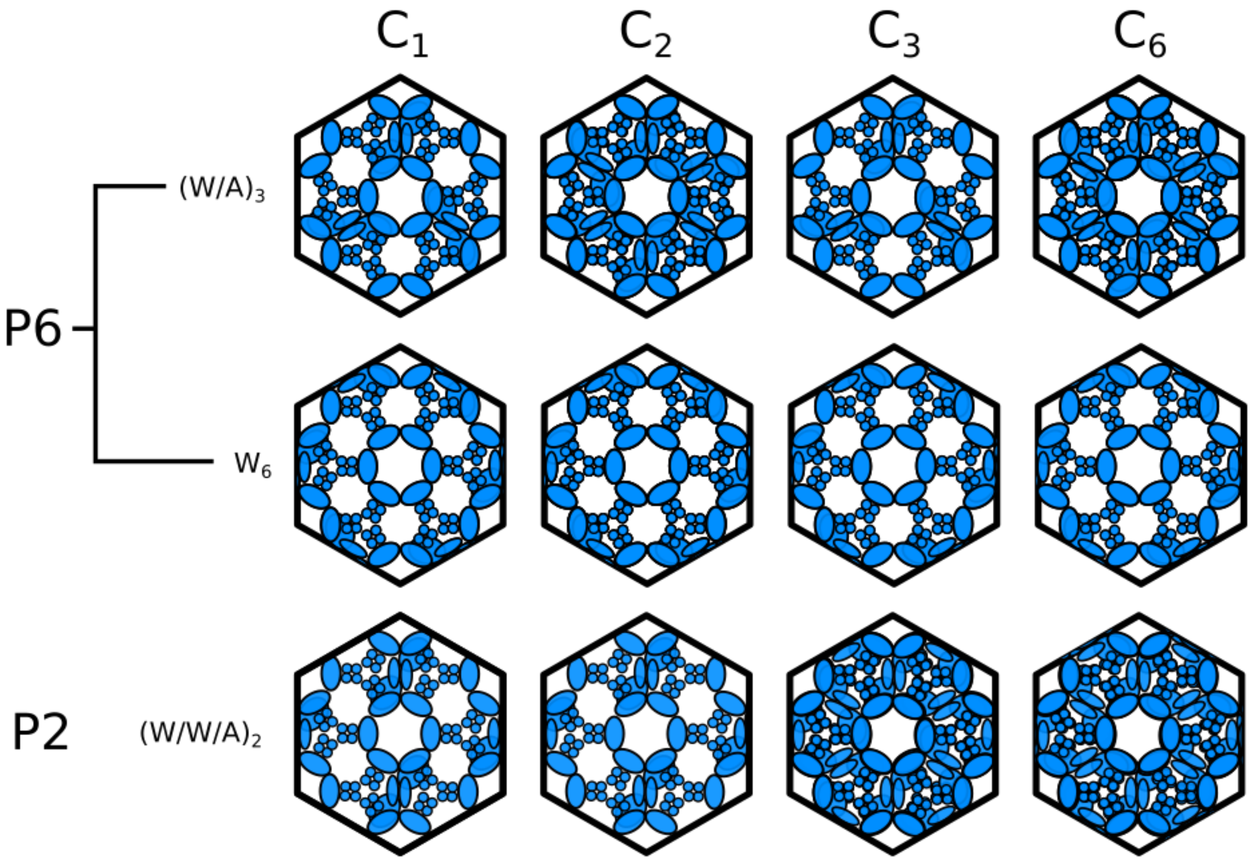


**Supplementary Figure 5. The effects of applying different symmetries to reconstructions centered at the center of six receptor trimers-of-dimers in both p6 and p2 architectures.**

**Movie 1. Dynamic exploration of a tomogram on a curved surface in the region of the chemosensory array.** Density from a WM4196 tomogram is projected onto a curved surface in the tomogram at the level of the chemosensory array inside the minicell. The region of density projected is shifted along the normal to the surface of each triangle in the mesh to explore the tomogram at different levels of the array architecture.

**Movie 2. A comparison of models of p6 and p2 chemosensory array architectures.** The models of both p6 and p2 symmetric array architectures are visualised as slices at different levels of the structure to highlight similarities and differences at various positions in the extended architecture. The arrow on the model of the chemosensory array on the left hand side indicates the position of the slice through the model of the extended architectures displayed on the right hand side.
